# Cortico-subcortical dynamics in primate working memory

**DOI:** 10.1101/2024.07.24.605013

**Authors:** Constantin von Nicolai, Markus Siegel

## Abstract

Working memory is essential for cognition, facilitating the temporary maintenance and manipulation of information to produce goal-directed behavior. While both cortical and subcortical structures are involved, their precise roles and interactions are not fully understood. To investigate this, we simultaneously recorded neural activity from the frontal and parietal cortex, higher-order thalamic nuclei, and core basal ganglia structures during color and spatial working memory tasks in non-human primates. We found widespread yet differential encoding of color and spatial information, marked by area-specific temporal dynamics and modulation according to task demands. Both cortical and subcortical information increased towards working memory-dependent actions, suggesting a task-specific reloading of information. Directed interactions between cortical and subcortical regions were extensive and reciprocal, with dominant directions of information flow, especially from frontoparietal areas. These interactions were dynamically modulated and partially task specific. Our findings provide comprehensive insights into the large-scale circuit dynamics underlying primate working memory and suggest that flexible goal-directed behavior relies on the selective processing of task-relevant memory information within information-specific cortico-subcortical networks.

## Introduction

Goal-directed behavior relies heavily on working memory, the capacity to retain sensory or internal information and use it to act according to current rules and task demands^1–4^. Working memory is a dynamic process with multiple stages from memory encoding to action that involves numerous interacting neural elements. Despite extensive research, the mechanisms by which the brain supports working memory are still not fully understood. This gap in understanding is partly due to the fragmented study of both the dynamics of working memory and the neural elements that jointly support it.

Regarding the dynamics of working memory, many studies have concentrated on mnemonic aspects such as memory encoding and maintenance, often overlooking memory utilization and the processes connecting these stages. While recent research has begun to address this gap^5^, invasive studies examining the neural basis of working memory from encoding to action are still rare.

In terms of neural elements, pioneering work has explored the neural correlates of working memory in various cortical and subcortical regions. Prefrontal and parietal cortical areas are known to play crucial roles in working memory^6–10^. Additionally, subcortical structures, particularly higher-order thalamic areas and basal ganglia regions, also likely contribute to working memory^11–17^. However, their fragmented study prevented a direct quantitative characterization how memory information is distributed across the cortico-subcortical network.

Furthermore, working memory, like other cognitive functions, fundamentally depends on interactions between multiple brain regions^18–22^. Despite this widely accepted view, with a few notable exceptions^21,23,24^, electrophysiological studies of working memory in primates have been conducted using recordings from one or at most two regions at a time. Studies using simultaneous recordings from cortical and subcortical areas are even more scarce. Consequently, due to the lack of simultaneous recordings, the interactions between areas that support working memory remain largely unknown.

In sum, three central issues and related questions regarding the neural basis of working memory persist. First, what are the dynamics of memory information from encoding to utilization? More specifically, is working memory information reloaded towards its use? If so, in which regions and is such an enhancement of information task specific? Second, what specific roles do key cortical and subcortical areas play? How is working memory information quantitatively distributed across the cortico-subcortical network? And, to what extent do higher-order thalamic nuclei and basal ganglia support other working memory features than spatial memory? Third, what is the structure of cortico-cortical and cortico-subcortical interactions supporting working memory? In particular, what is the dominant direction of interactions within the frontoparietal network or between cortical and subcortical regions? And how are network interactions modulated during working memory from encoding to action?

To answer these questions, we simultaneously recorded neural activity from prefrontal, parietal, thalamic, and basal ganglia regions in macaque monkeys during color and spatial working memory tasks. These large-scale recordings provided us with the unique opportunity to directly compare mnemonic information in all these structures and to quantify their interactions.

We found that color and spatial memory information are widely distributed across all investigated cortical and subcortical structures and that they exhibit dynamics that vary by area and in a task-dependent manner. Notably, the utilization of working-memory information for action is preceded by the reloading of task-relevant information in information-specific cortico-subcortical networks. Reciprocal interactions between cortical and subcortical areas are ubiquitous, dynamic, and partially task-specific, with interactions again being particularly strong during memory use for action. Overall, cortical areas LIP and FEF appear to serve a coordinating role in this network by driving other regions.

## Results

### Task, recordings, and behavioral results

We trained two macaque monkeys in a working memory task in which the animals had to memorize either the color or the position of a peripheral cue over a delay period of 1.25 s and to respond with a saccade either to the matching one of two colored peripheral targets or to the position of the preceding cue (color and spatial task, respectively; Fig. 1A and Methods). Importantly, the visual input was the same for both tasks until the beginning of the response period. Furthermore, in the spatial task, animals could in principle plan the required upcoming eye movement to the target to be memorized from the cue presentation onwards, whereas in the color task they could not. The conditions were presented in blocks without explicit cueing of the current task rule, i.e., the task at hand was implicit. Both animals performed well on both tasks, with percentage correct performance being 83% (monkey V) and 80% (monkey E) for the color task and 94% (monkey V) and 92% (monkey E) for the spatial working memory task.

**Figure 1.**
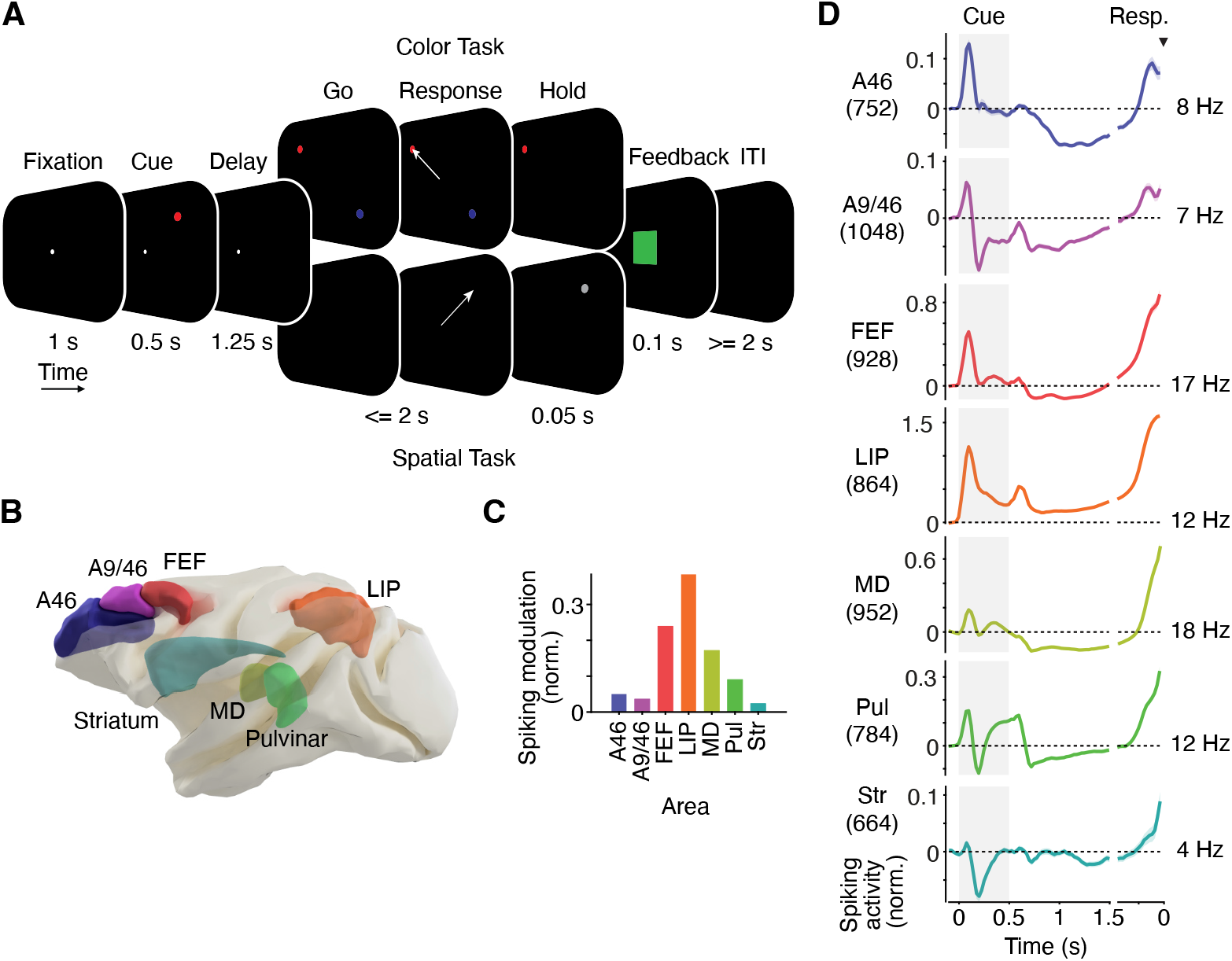
Behavioral tasks, target regions and spiking activity. (**A**) Behavioral tasks. Monkeys performed a color and a spatial working memory task in block-wise manner. For both tasks, a colored cue (blue or red) was presented at one of four spatial locations (90, 180, 270, 360 degrees) for 0.5 s. Following a 1.25 s delay and fixation spot offset, monkeys either had to saccade to the matching color target presented together with the non-matching target in a random spatial configuration across both main diagonals (color task) or towards the remembered stimulus location (spatial task). For correct responses, a liquid reward was delivered, and a green square was flashed. (**B**) 3D-rendering of the brain and target regions for recordings. (**C**) Spiking activity modulation quantified as the coefficient of variation across trial time (standard deviation relative to the mean). (**D**) Average normalized spiking activity during both tasks. Shaded regions denote SEM. Numbers in brackets and on the right indicate number of electrodes and baseline firing rates, respectively.

Once the animals had mastered the task, we implanted recording chambers over large parts of one hemisphere (Supplementary Figure S1). This provided simultaneous access to seven frontal (A46, A9/46, frontal eye field - FEF), parietal (lateral intraparietal area - LIP), higher order thalamic (mediodorsal thalamus - MD and Pulvinar) and basal ganglia (Striatum) brain regions (Figure 1B, Supplementary Figure S1 and Methods). We simultaneously recorded extra-cellular spiking activity from these regions using multi-contact microelectrodes.

### Dynamics of spiking activity

We found that spiking activity was substantially modulated by task events in all recorded areas (Figure 1C and D). This temporal modulation was strongest in LIP and weakest in the Striatum and prefrontal areas (Figure 1C). The temporal profile of spiking dynamics was generally similar across brain regions. Spiking transiently increased following stimulus onset and then decreased towards the memory delay. Interestingly, during the delay, spiking remained elevated only in area LIP, but dropped below baseline in all other areas. Towards the response period, spiking significantly increased again in all regions.

### Dynamics of memory information

To address the first two questions raised above, about the dynamics of working memory information and how it unfolds in cortical and subcortical areas, we quantified the information about critical task variables encoded by spiking activity. Specifically, for each time point throughout the task, we determined the variance in spiking that could be explained by the color and spatial location of the memory cue across trials (color and spatial information) Importantly, our approach ensured that information about each task variable was quantified independently from all other variables (see Methods).

Figure 2 shows the time course of neural information during correct trials about the color and spatial position of the cue from the prestimulus baseline through the memory delay to response onset (Figure 2A) as well as the average information and corresponding statistics during three successive time windows: cue presentation (0-500 ms post cue onset), early memory delay (250-750 ms post cue offset), and late delay before the response (500-0 ms before response onset) (Figure 2B). To quantify and statistically assess the temporal dynamics of information, we computed differences of information between successive task periods (Figure 2C).

**Figure 2.**
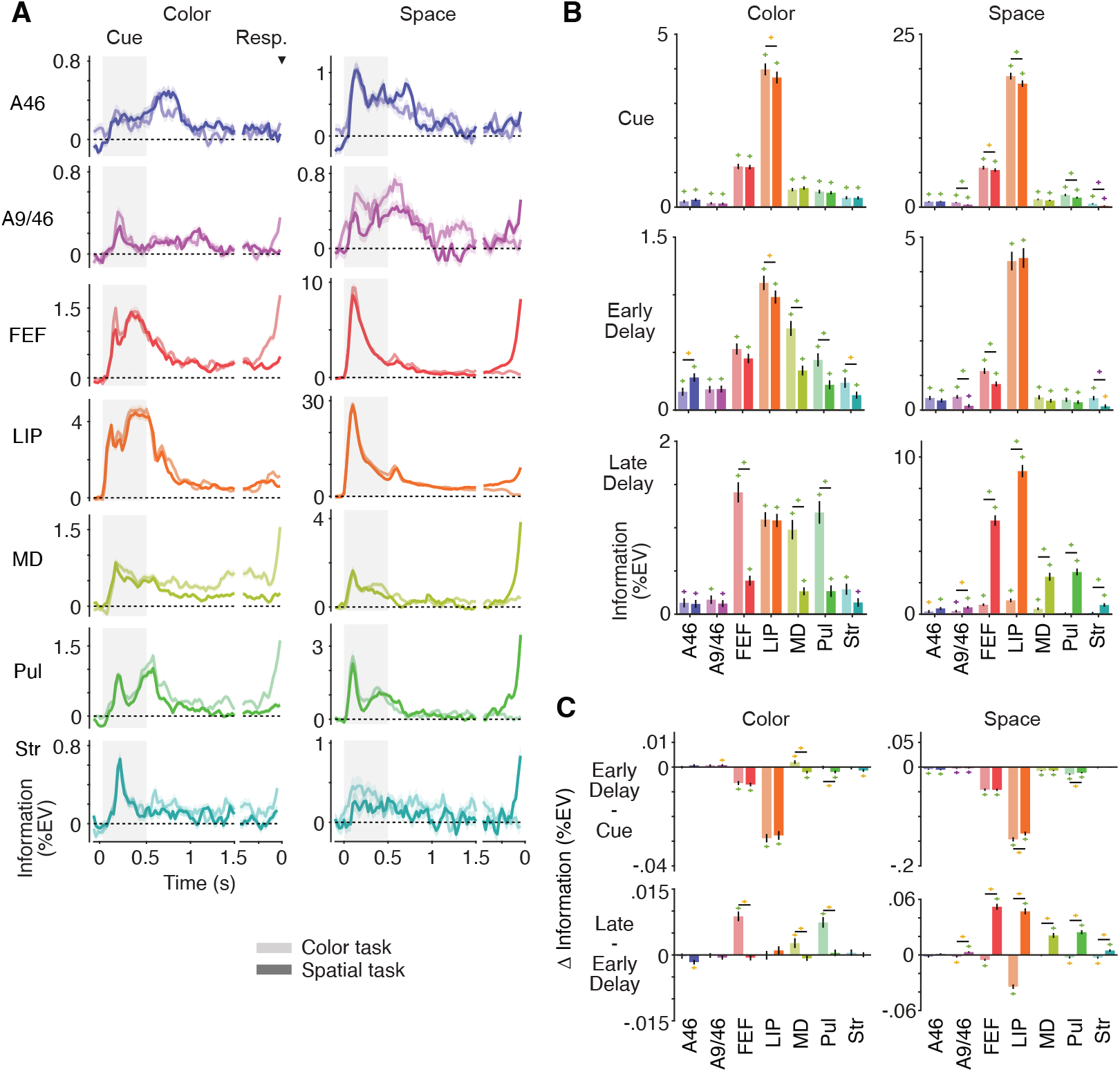
Dynamics of memory information. (**A**) Time courses of information about the color and spatial position of the memory cue during correct trials of the color (light) and spatial (dark) working memory tasks. (**B**) Fixed-window statistics on color and spatial information during 3 task periods. (**C**) Fixed-window difference statistics on temporal changes of color and spatial information. Shaded regions and error bars denote SEM. Colored crosses denote significance (yellow, purple, green: p < .05, .01, .001, respectively; paired and unpaired t-tests, FDR-corrected).

We found that, following a variable rise upon cue onset, during cue presentation, color and spatial information was present in all recorded cortical and subcortical areas (Figure 2A and 2B, first row). Following cue offset, information dropped again in most areas, while notably MD and A9/46 showed stronger color information during the early delay of the color and the spatial task, respectively, as compared to the cue period (Figure 2C, first row). Throughout the entire memory delay, all cortical and subcortical areas significantly encoded color or spatial information with variable strength (Figure 2B, middle and bottom row). Thus, despite the wide-spread suppression of spiking activity, during the memory delay (Figure 1D), task-relevant memory information was broadly distributed across cortical and subcortical structures.

Also, in the late delay interval, when the memory-based response was imminent, color or spatial information was significant in all recorded cortical and subcortical regions (Figures 2B, bottom row). Moreover, there was a marked increase of memory information towards the response in distinct sets of regions depending on the task and information at hand (Figure 2A and C).

In sum, during cue presentation and the following memory interval, visual color and spatial information was broadly encoded across parietal (LIP), frontal (FEF) and prefrontal cortex (A46, A9/46), the thalamus (MD, Pulvinar) and Striatum. Furthermore, in this network, working memory information was prominently reloaded towards the memory-guided behavioral response.

### Task dependence of memory information

How did the behavioral task that the animals had to perform (report either memorized color or space) affect the encoding of visual information? We found that particularly the reloading of memory information towards the response was significantly modulated by the task at hand (Figure 3). During cue presentation and the early delay, both color and spatial information were stronger almost exclusively in the color task (Figure 3A and B). This changed markedly during the late delay when utilization of mnemonic information for the response was imminent. At this point, not only did color and spatial information increase relative to the early delay (Figure 2C bottom row), but color and spatial information were specifically enhanced in the task for which the corresponding information was behaviorally relevant. In the late delay, color information was significantly stronger when color had to be reported and spatial information was significantly stronger when space had to be reported (Figure 3C). This task-dependence was due to a selective reloading of information. The irrelevant information showed no significant enhancement towards the response (Figure 2C). This task-dependent modulation towards the response was significant in distinct sets of cortical and subcortical regions depending on the task and information at hand (Figure 3C).

**Figure 3.**
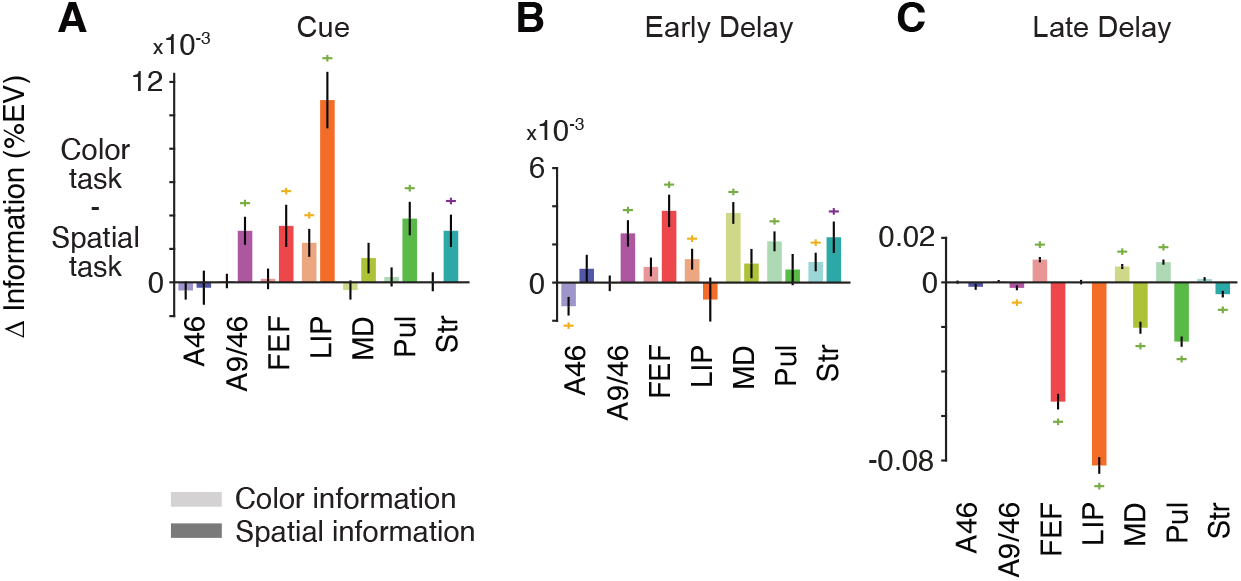
Task dependence of memory information. (**A**) Fixed-window statistic on task-related differences (color task - spatial task) of color and spatial information in the cue interval. Positive and negative values indicate more information in the color and spatial task, respectively. (**B**) Task-related differences in the early delay interval. (**C**) Task-related differences in the late delay interval. Error bars denote SEM. Colored crosses denote significance (yellow, purple, green: p<.05, .01, .001, respectively; unpaired t-tests, FDR-corrected).

Despite this abundance of the late task-dependent modulation, the time-course of this effect differed between regions. MD appeared to be the first structure to exhibit a clear task-specific enhancement of color information already during the early delay period (Figure 2A and 3B). A weaker early enhancement during the delay was also seen in in the Pulvinar, Striatum and LIP (Figure 2A and 3B).

In sum, we found that encoding of visual information was modulated by the task at hand. Towards the behavioral response, specifically the memory information that was relevant for the task at hand was enhanced in specific cortico-subcortical networks.

### Interactions between space and color

The above analysis quantified information about color and space as independent variables. In another set of analysis, we investigated the dynamics of information about the interaction of color and space, i.e. neural activity that encoded a specific color at a specific spatial location (Figure 4A and B). Do only visuomotor areas such as LIP or FEF carry retinotopically specific memory information? Or is memory information also spatially specific in other cortical or subcortical regions? We found evidence for the latter.

**Figure 4.**
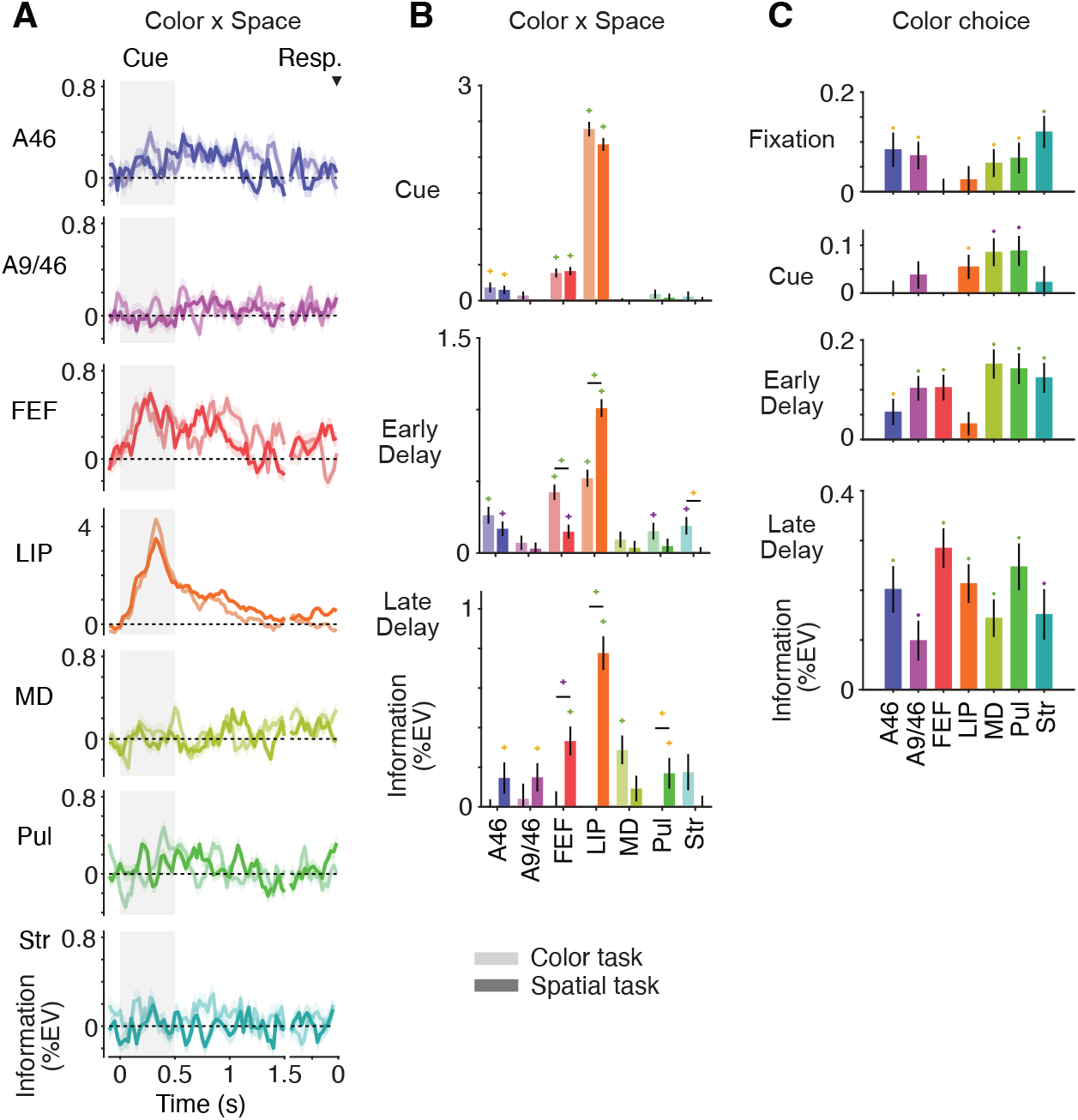
Interaction between color and spatial information and color choice information. (**A**) Time courses of information about the interaction between the color and spatial position of the memory cue during color (light) and spatial (dark) working memory tasks. (**B**) Fixed-window statistics on information about the interaction between color and space during 3 task periods. (**C**) Fixed-window statistics on chosen color information during four periods of the color working memory task. Shaded regions and error bars denote SEM. Colored crosses denote significance (yellow, purple, green: p<.05, .01, .001, respectively; paired and unpaired t-tests, FDR-corrected).

Throughout the trial spatially specific color information was strongest in LIP and FEF. However, during cue presentation, also A46 exhibited a significant interaction effect (Figure 4A and B). In addition, in the early delay the interaction between color and space was also significant in the color task in the Pulvinar and Striatum whereas in the spatial task, it was significant only towards the response in all areas except for MD and the Striatum. Thus, although spatially specific color information was most prominent in visuo-motor areas LIP and FEF it was also broadly distributed across the cortico-subcortical network.

### Choice information

During the color task, animals made errors, i.e. they did not always choose the cued color (17% for monkey V; 20% for monkey E). This allowed us to investigate neural information about the animals’ color choice independent of the sensory input (Figure 4C). Importantly, as the color response targets were spatially randomized, color choice information was also independent of the spatial motor response.

We found significant choice information, i.e. neural activity that predicted the animals’ color choice independent of the sensory cue and motor response, during all trial intervals including the fixation baseline. Choice information was generally strongest in the late delay before the behavioral response, but the distribution of choice information across brain regions changed markedly throughout the trial. During the pre-cue baseline, choice predictive information was strongest in prefrontal (A46, A9/46) and subcortical (MD, Pulvinar, Striatum) regions and not significant in LIP and FEF (Figure 4A). During cue presentation, choice information was strongest in the thalamus (MD, Pulvinar) and significant in LIP. During the delay, choice information increased such that during the late delay before response onset, choice predictive information was broadly distributed and significant across all cortical and subcortical areas. In sum, choice information showed a dynamic remapping from being confined to associative and subcortical regions before the sensory input to a broad distribution across the entire sensorimotor pathway and subcortical regions towards the choice report.

In sum, we found widespread dynamic correlates of working memory for action in cortical and subcortical regions. Memory information was broadly distributed across the cortex, thalamic nuclei and basal ganglia. Task-relevant memory information was enhanced when memory-based action was impending in distinct but overlapping sets of cortical and subcortical areas with thalamic nuclei (MD and Pulvinar) showing an early task-specific enhancement of memory information. Furthermore, color choice predictive information was already present in subcortical and associative cortical regions before the memory cue and was then widely distributed across all cortical and subcortical regions before the choice report.

### Directed interactions between cortical and subcortical areas

We next asked how cortical and subcortical structures interacted during the working memory task. To investigate this, we computed Granger causality between spiking activity from all simultaneously recorded areas (Figure 5). We applied strict statistical criteria to rule out spurious interactions (see Methods). Specifically, we required a given area pair to fulfill three requirements: first, significant shuffle-corrected Granger causality; second, significant interareal asymmetry of Granger causality, indicating a dominant direction of information flow; and third, a reversal of the interareal asymmetry for time-reversed Granger causality^25,26^. In addition to computing Granger causality between pairs of areas, we also determined each areas’ average interaction with all other structures.

**Figure 5.**
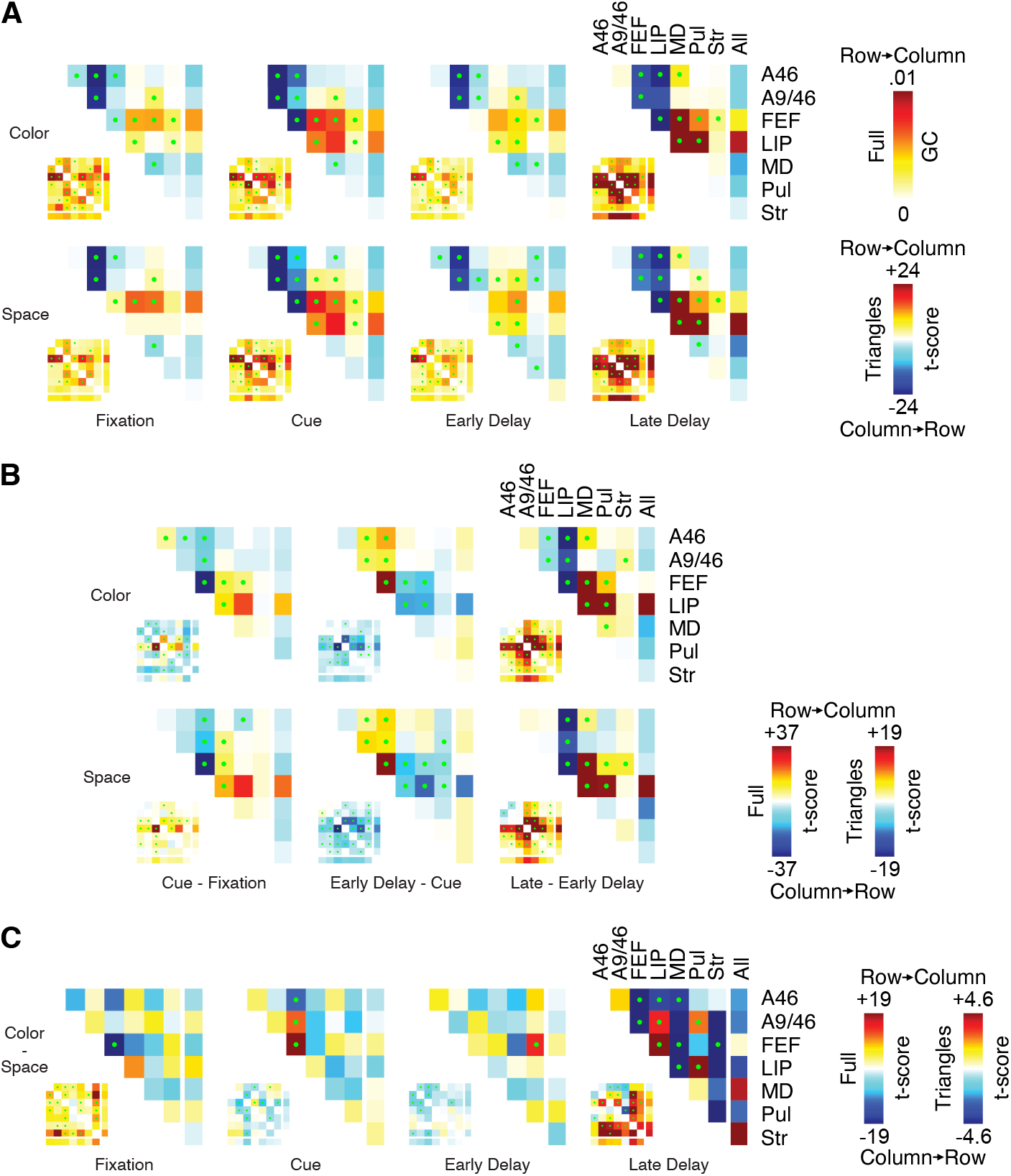
Directed cortico-subcortical interactions. Granger causality (full matrices) and asymmetry of Granger causality (triangular matrices) between spiking activity in pairs of brain regions. Full matrices show interactions in both directions. Triangular matrices show the asymmetry of interactions, i.e. the difference between both directions. The last row and/or column depict the corresponding marginal interactions. (**A**) Full directed interactions and directional asymmetries during four periods of the working memory tasks. Asymmetries were computed by contrasting interactions of both directions of a given area pair. Asymmetries are shown as t-scores across electrode pairs. For full matrices, rows and columns correspond to sources and targets of interactions, respectively. For triangular matrices, asymmetries are from row to column for positive values and reversed for negative values. (**B**) Temporal differences of interactions across three successive task periods. Plots show t-scores of window differences across electrode pairs. (**C**) Differences of directed interactions between color and spatial working memory tasks. Plots show t-scores of task differences (color - space) across electrode pairs. Green dots denote area pairs with significant Granger causality in either direction after shuffle correction (p<.05; FDR-corrected) and a significant asymmetry (p<.05; FDR-corrected) that reverses for time-reversed data.

We found widespread, reciprocal interactions between areas in both tasks (Figure 5A, small full matrices). For each pair of areas, we subtracted both directions of interactions to quantify the asymmetry or dominant direction of information flow. FEF and LIP were by far the strongest senders, interacting with almost all other areas in a driving manner. In contrast, all other structures were by and large net receivers, including thalamic areas MD and Pulvinar as well as the Striatum (Figure 5A, large triangular matrices). Interestingly, interactions between prefrontal areas A46 and A9/46 were very rare. They were net receivers of information from FEF and LIP but had a net driving influence on MD and Pulvinar. Most importantly, directed interactions were strongest during the late delay when memory information was needed for response generation.

As with information encoding, we next tested whether the structure of directed interareal interactions was dynamic (Figure 5B). We computed differences in Granger causality between the same consecutive temporal window pairs as above. In addition, given the substantial interactions already during the fixation baseline, we also quantified the change in Granger causality from fixation to memory cue presentation. Directional coupling increased with cue presentation, particularly in area pairs with LIP and FEF in both tasks (Figure 5B, small full matrices). Also, the net drive exerted by FEF and LIP on thalamic areas MD and Pulvinar increased with cue presentation (Figure 5B left, large triangular matrices). Interestingly, some pairs of areas in the color task also showed reduced interaction during cue presentation. From cue presentation to early delay, interactions mostly decreased, and the net driving influence of FEF and LIP on prefrontal and thalamic areas also decreased. From early to late delay, interactions became ubiquitously stronger and the net driving influence of FEF and LIP on MD and Pulvinar increased again.

Finally, we examined whether the structure of directed interactions was task-specific (Figure 5C). During fixation, raw Granger causality (small full matrices) was stronger in the color task than in the spatial task, whereas directional asymmetries (large triangular matrices) were hardly task-specific, except for a stronger drive from LIP to FEF in the spatial task. Interestingly, during cue presentation, the task-specificity of directional asymmetries was limited to interactions involving the largely visual area LIP. During the early delay, raw interactions were generally stronger in the spatial task, and this task-specificity was particularly pronounced in area pairs involving frontal areas A46, A9/46, and FEF.

However, there were hardly any task-specific directional asymmetries during this period. This changed markedly during the late delay, when the animals prepared for action based on working memory. Towards the behavioral response, many pairs of areas showed task-specific interactions and directional asymmetries. In particular, MD and Pulvinar showed stronger sending relationships with many areas during the color and spatial tasks, respectively. Overall, frontal, parietal, and thalamic structures showed substantial task-specific modulation before the response.

In summary, we found ubiquitous cortico-cortical and cortico-subcortical directed interactions that were clearly structured and dynamically modulated during the working memory tasks. Frontoparietal areas FEF and LIP played major driving roles during all task phases, whereas prefrontal and subcortical areas were predominantly receivers of information. Overall, directed interaction increased significantly during preparation for action. During this period, and especially between frontoparietal and thalamic regions, directed interactions were also most task specific.

## Discussion

Our study offers new and comprehensive insights into the large-scale circuit mechanisms that underlie working memory. We leveraged an advanced experimental approach with simultaneous large-scale recordings from several cortical and subcortical regions of the primate brain during a flexible working memory task. Our approach revealed that color and spatial memory information is pervasive in prefrontal, parietal, thalamic, and basal ganglia structures, exhibiting area-specific and task-dependent dynamics. We observed that the utilization of working memory information for action is preceded by the reloading of task-relevant information in overlapping but distinct cortical and subcortical regions. Additionally, reciprocal interactions between these areas were widespread, dynamic, and partially task-specific, with the strongest interactions occurring during memory use for action. Specific cortical areas, particularly LIP and FEF, played a dominant role in driving activity in other regions.

### Neural dynamics of working memory processing

Our findings enhance the limited understanding of the neural dynamics involved in working memory-dependent action. It has been suggested that internal selective attention functions are crucial for both mnemonic and action-related aspects of working memory^4^. This involves enhancing relevant over irrelevant working memory representations for action^27^. Our observation of task-specific enhancement of either color or spatial information in terms of a reloading of information towards the response supports this idea. Furthermore, our results indicate that the cortico-subcortical networks prioritizing task-relevant information are specific to the type of information. While all areas supported information encoding and maintenance, reloading of task-relevant information towards the response was task- and information-specific.

During the late delay, color information increased in FEF, MD, and Pulvinar in the color task, but not in the spatial task. Conversely, spatial information increased only in the late delay of the spatial task in A9/46, FEF, LIP, MD, Pulvinar, and the Striatum. Memory information notably increased late during the delay in both tasks, suggesting an action-bound and task-specific selection of memory information in large-scale cortical-subcortical networks^4,22^.

Our study also revealed intriguing dynamics of stimulus-independent neural activity predicting the animals’ color choice. Choice information was present during the fixation baseline in many areas, excluding FEF and LIP, but became broadly distributed across all cortical and subcortical areas towards the response. These findings align with the distributed encoding of choices observed in cortical and subcortical areas in rodents^28^ and primates^29^. Moreover, the choice signal dynamics demonstrated in the present study suggest that interactions between rostral prefrontal regions, along with associated higher-order thalamic and basal ganglia regions, may influence activity in structures more dedicated to memory-dependent action preparation.

### Cortico-subcortical interactions during working memory

Our large-scale simultaneous recordings allowed us to, for the first time, characterize directed interactions between key cortical and subcortical regions during working memory in the primate brain. Critically, we directly quantified interactions at the level of spiking activity, which avoids potential interpretational ambiguities at the level of populations signals such as e.g. local field potentials. We identified strong reciprocal interactions, with many area pairs showing directional asymmetry, indicating a dominant direction of interaction. These interactions were dynamic, with a prominent enhancement towards the behavioral response. They were also task-specific, with distinct connections being enhanced during either color or spatial working memory.

Overall, LIP and FEF were primarily net senders of information, while prefrontal and subcortical regions predominantly received information. These findings align with previous research that demonstrated a stronger driving influence of the parietal cortex on the prefrontal cortex during working memory^21^, supporting the idea that frontoparietal network interactions are crucial for working memory processing^21,30–32^.

Our approach also enabled the examination of interactions between frontoparietal areas and their primary higher-order thalamic partners. The MD and Pulvinar thalamic nuclei are known to facilitate interactions between cortical areas during cognitive tasks^13,14,33–39^. While previous studies have shown that the thalamus, such as MD on the prefrontal cortex^13,14,38,39^ or the Pulvinar on parietal areas^35,36^, can exert a strong driving influence, this seems to depend on the specific task and stage examined. Reciprocal interactions appear to be the norm rather than the exception^33^. Indeed, we observed significant reciprocal interactions between cortical and thalamic areas, with cortical drive often exceeding thalamic drive in both working memory tasks. Additionally, thalamic areas demonstrated substantial task-specific information processing, sometimes even earlier than cortical areas, such as MD during the color task. These results highlight the important role of corticothalamic interactions in working memory and suggest an early, task-dependent selection of memory information in subcortical structures, independent of dominant cortical influence.

### Specific roles of cortical and subcortical regions in working memory

Our study offers novel insights into the specific roles of various cortical and subcortical regions in working memory, uncovering task-dependent dynamics and emphasizing their contributions to processing and utilizing memory information for action. Notably, the simultaneous recordings during the same behavior allowed us to directly quantify the distribution of memory information across a broad cortico-subcortical network.

We did not observe a particularly prominent role of prefrontal areas A46 and A9/46 in working memory information coding and directional coupling. At first glance, this may seem contrary to their known involvement in working memory^40,41^. However, many prior investigations on prefrontal mechanisms featured recordings from caudal PFC regions like FEF (area 8Ad). Indeed, we found that FEF exhibited strong task-dependent memory encoding and exerted robust influences on all other areas except LIP. This confirms previous studies suggesting that FEF, as a mixed visuomotor structure, is crucial for preparing eye movements^42^ and is a key component of attention and working memory networks, at least in visual tasks requiring eye movements^43,44^. Our results suggest that FEF plays a central role in transforming memory information into eye-movement responses through interactions with cortical (LIP) and subcortical (MD, Pulvinar) partners. Future studies are required to test if more demanding tasks rely more on coordinating activity in rostral frontal areas like A46 and A9/46^44,45^.

Parietal area LIP exhibited the strongest color and spatial information coding during cue presentation, which aligns with its role in visual and spatial cognition^46^. Unlike FEF or thalamic areas, LIP’s color information was not enhanced during the late delay when memory information was used for action. However, LIP was the strongest source of directed interactions during both cue presentation and the late delay of both tasks, even compared to FEF, and had a net driving influence on thalamic areas throughout the delay. Our results suggest that LIP significantly contributes to information maintenance during mnemonic delays and provides essential input to areas like FEF for generating behavioral responses^44,47,48^.

Thalamic areas, particularly MD, were first to show task-dependent differentiation of cue information coding during the memory delay and towards the behavioral response. MD has long been associated with working memory processing, similar to the prefrontal cortex, its main input and output region^10,11,13,49,50^ (but see ^3,51–53^). Our findings align with previous evidence that MD supports sensory-to-motor transformations and coordinates frontal cortex computations in a task-dependent manner^12–14,33,49,54–56^. Prior studies on MD have primarily focused on spatial working memory tasks^13,57–59^. Our results indicate that MD supports working memory more generally by maintaining and reloading also non-spatial memory information needed for action.

The Pulvinar, unlike MD, has been primarily linked to visual sensory and attentional processing. However, it is a higher-order thalamic nucleus with extensive connections to occipital, temporal, parietal, and frontal cortex areas, including FEF and dorsolateral PFC^60^. This suggests that in particular the dorsomedial Pulvinar studied here is involved in cognitive processes like working memory^61,62^. Indeed, we found that the Pulvinar exhibits working memory-related coding and connectivity similar to MD, with differences in interaction partners during specific task stages. Given their overlapping connections, this can be expected, but the Pulvinar’s broader connections might lead to different dynamics. Future studies should clarify how MD and the dorsomedial Pulvinar support cognition together. In any case, our results emphasize the important roles of higher-order thalamic nuclei and thalamocortical loops in cognition^17,52,54,63^.

The Striatum, a significant site of dopaminergic influence^15,16,64^, has been linked to working memory processing^65^ because it is the primary input structure of the basal ganglia for cortical projections, establishing cortico-basal ganglia-thalamocortical loops^18,66^. Early studies showed striatal activity modulation during different stages of working memory tasks^67^, pointing to a possible role as a subcortical gate-keeper^65,68^. Indeed, recent fMRI studies found evidence for corticostriatal output gating during working memory tasks^69^. The Striatum was the least prominent sender and receiver in the interaction analyses, possibly due to challenges in targeting interconnected sites in this large structure or the nature of the tasks used^70,71^. However, we found significant cue-related color and spatial memory information in the Striatum, which suggests its contribution to both spatial and non-spatial working memory for action in the primate brain.

## Conclusion

Our findings indicate that task-relevant information is widely distributed across key cortical and associated subcortical areas during color and spatial working memory tasks, with dynamics that vary by task and area, primarily bound to action. Both the frontoparietal cortex and higher-order thalamic regions play significant roles in working memory processing with frontoparietal areas FEF and LIP serving central, coordinating functions. Extensive interactions between cortical and subcortical areas reflect task engagement and adapt to task demands, particularly during memory for action. These insights clarify how working memory integrates sensory information with its use for action bridging the perception-action cycle^4,22,72^.

## Acknowledgements

We thank Peter Dicke for technical advise, assistance and helpful discussions. This research was supported by the European Research Council (ERC; https://erc.europa.eu/) StG 335880 (M.S) and CoG 864491 (M.S), Deutsche Forschungsgemeinschaft (DFG; German Research Foundation; https://www.dfg.de/) project SI1332/3-1 (M.S.) and project 276693517 (SFB 1233) (M.S.) and the Centre for Integrative Neuroscience (DFG, EXC 307) (M.S.). The authors acknowledge support by the state of Baden-Württemberg through bwHPC, by the German Research Foundation (DFG) through grant no INST 39/963-1 FUGG (bwForCluster NEMO), and by the Open Access Publishing Fund of the University of Tübingen. The funders had no role in study design, data collection and analysis, decision to publish, or preparation of the manuscript.

## Author contributions

CvN: Conceptualization, Methodology, Investigation, Formal analysis, Visualization, Writing – original draft, Writing – Review & Editing MS: Conceptualization, Methodology, Investigation, Supervision, Resources, Project administration, Funding acquisition, Writing – original draft, Writing – Review & Editing

## Data availability statement

Data and code to reproduce all the reported results are available from the lead author upon reasonable request.

## Methods

### Subjects and behavioral task

All procedures involving non-human primates were carried out in agreement with national and international law, rules and regulations on the humane care and treatment of research animals and were approved by local authorities.

We trained 2 macaque monkeys (*Macaca mulatta*) in two tasks that probed color and spatial working memory, respectively (Figure 1A). Importantly, both tasks had identical, fixation baseline, cue presentation and memory delay intervals. Tasks were presented in a block-wise manner (at least 64 correct trials per block), i.e., the current task rule was implicit. Animals initiated a trial by acquiring and holding fixation of a central white spot (diameter: 0.1 º visual angle) presented on a black background for 1 s. Following fixation, a blue or red cue stimulus (diameter: 1 º) was presented for 0.5 s at 45, 135, 225, or 315 º circular angle and 6 º eccentricity. Depending on the current task, animals had to memorize either the color or the location (color and spatial task, respectively) of the stimulus over a delay of 1.25 s during which they had to maintain central fixation. Following the delay and the offset of the fixation point, in the color task, a pair of peripheral targets colored blue and red was presented at two randomly chosen, opposing (180 º circular distance), possible prior stimulus locations. In the spatial task, fixation point offset was followed by a blank screen. Animals had to make a saccade either to the target that matched the previously cued color (color task) or to the previously cued location (spatial task). Thus, in the spatial task, animals could prepare the motor response during the delay whereas in the color task, they could not. In both tasks, monkeys had to hold their gaze at the correct target location for at least 50 ms. Then, for correct trials, a green central square (edge length: 6 º) was shown and animals received a liquid reward. An incorrect response was followed by a 100 ms flash of a grey square of the same size. A trial was aborted if monkeys did not keep their gaze within a 1.5 º radius around the fixation spot before the end of the delay period. Both correct and error trials were followed by an inter-trial period of at least 2 s.

Behavioral task control was accomplished with the Monkeylogic toolbox^73^ for MATLAB (The Mathworks, Natick, MA, USA). Stimuli were presented using CRT monitors with a vertical refresh rate of 100 Hz. Monkeys sat in a primate chair (Crist Instruments) at a viewing distance of 42 cm from the screen and their heads were immobilized via titanium headposts throughout the course of behavioral training and recording sessions. Gaze control was accomplished with infrared eyetracking systems (View-Point EyeTracker, Arrington Research Inc., Scottsdale, Arizona, USA; and Eyelink 1000, SR Research, London, Ontario, Canada) at a sampling rate of 220 and 1000 Hz; we used the Eyelink system for neural recording sessions. We measured the luminance of monitor background, fixation spot, cue and target stimulus, and feedback square colors and equated their brightness for behavioral training and recording sessions.

### Neural recordings

Electrodes were lowered into and removed from the brain anew on each recording day. We used custom-designed, 3D-printed grids for placement of electrode drives and precise, reliable targeting of cortical and subcortical structures (Fig. 1B and Figure S1). We recorded broadband neural activity using linear, multi-contact Plexon (Plexon Inc.; Dallas, Texas, USA) or Neuronexus (NeuroNexus Technologies, Inc.; Ann Arbor, Michigan, USA) probes at a sampling rate of 24414 Hz with a TDT (Tucker-Davis Technologies; Alachua, Florida, USA) multichannel recording system. We used Plexon U- and V-probes with 24 contacts and vertical interelectrode distances of either 0.1, 0.15, 0.25, or 0.5 mm (single electrode configuration) or 0.15 mm vertical and 0.05 mm horizontal inter-electrode distance (stereotrode configuration); Plexon S-probes with 32 contacts and vertical interelectrode distances of 0.1 or 0.25 mm; and Neuronexus probes with 32 contacts and vertical inter-electrode distances of 0.1 mm.

### Area targeting

For chamber implantation and recording target planning, we resliced T1-weighted structural MR images of both animals such that the interaural zero point represented the origin of the recording space^74^ (Figure S1). We used a publicly available, standardized rhesus macaque brain atlas^75^ to perform successive, combined linear and non-linear coregistriations between this atlas, regions of interest (ROI) label maps derived from it and the individual MR images of the animals. This provided region label volumes that we combined with MR images for target planning, in addition to considering other standardized rhesus macaque brain atlases^74,76^ and various literature sources on the anatomical connectivity between our target regions. We custom-designed titanium chambers based on the individual animal’s skull surface above the desired target brain areas using CAD-software, with the construction space being equal to the interaural space used for target planning. Headpost and chamber implantations were performed in separate sessions under general anesthesia and aseptic conditions. For chamber implantations, a stereotaxic arm was used to position the chamber at its target location in interaural space. Following implantation, we measured of the exact position of the chamber in 3D-space and realigned the final position of the chamber with the individual monkey’s MRI images to allow for precise target planning using 3D-printed grids placed inside the chamber.

### Signal preprocessing

From the broadband recordings, we derived spiking signals (multi-unit activity; MUA) by 500 Hz high-pass filtering. For analyses of neural information and directed interactions of spiking activity, we made use of the continuous time course (envelope) of MUA, so-called analog multi-unit activity (AMUA), which provides a robust estimate of neural population activity. We computed AMUA by rectification, 250 Hz low-pass filtering and logarithmic scaling of the high-pass filtered data. To account for slow drifts and offsets of spiking activity over time we subtracted a median filtered AMUA signal (10 trials) prior to information and Granger causality analyses.

### Spiking information coding

We computed AMUA information about variables of interest by means of two separate n-way ANOVAs across trials for each recording site using *μ*^*2*^ as an unbiased measure of variance of spiking activity explained^77^. The first ANOVA served to compute information about cue color (blue vs. red), cue location (4 locations at 45, 135, 225, 315 º circular angle) and their interaction for correct trials. It included the factors cue color, cue location, response location and a running count of the respective task. We conditioned it on the factors task (color vs. spatial task) and block. The second ANOVA served to compute information about the color of the chosen target. It included the factors task, cue color, cue position, chosen target color and response location; we conditioned it on the factor block. For each ANOVA, block-wise estimates of information where averaged before further analyses. For information time courses, we used a sliding window approach (50 ms window, 25 ms steps). For fixed temporal windows, we used non-overlapping 250 ms windows. These windows were used for further statistical analyses.

### Granger causality analysis

We quantified broadband Granger causality between pairs of AMUA signals by integrating the frequency-domain estimates across the spectrum^78^. Frequency-domain estimates were based on Fourier-transforms computed for frequencies between 0 and 256 Hz in 4 Hz steps for 0.25 second segments of AMUA after multiplication with 3 discrete prolate spheroidal sequence (DPSS) tapers. To avoid erroneous inferences of statistical causality due to volume conduction effects, we computed Granger causality on trial-shuffled and on time-reversed AMUA signals^25^. We subtracted the trial-shuffled Granger causality from all raw Granger causality estimates.

### Statistics on neural information

We used one-sided t-statistics to evaluate the significance of explained variance against the null hypothesis of zero information in separate time windows spanning 500 or 250 ms each. For the fixation, stimulus presentation, and early delay periods, we averaged omega square values of two adjacent 250 ms segments to comply with the 250 ms window of the late delay period. For time-window differences, we subtracted omega square values of the corresponding condition-specific windows and, for normalization, divided the difference by their sum. Similarly, for condition differences, we subtracted omega square values of condition-specific windows and normalized by their sum. We used two-sided t-statistics to evaluate significance of window differences from 0 of individual conditions, condition differences and differences between condition-specific window-differences. Alpha levels of all t-tests were set to p < 0.05. All p-values were FDR-corrected according to the method proposed by Benjamini and Hochberg^79^ to account for multiple hypotheses testing.

### Statistics on Granger causality

We applied several criteria to Granger causality to be taken as reflecting true interaction between a given pair of areas. First, Granger causality from either direction A to B or B to A had to be significantly non-zero (one-sided t-test across electrode pairs, p < 0.05 FDR corrected) after subtracting a surrogate estimate obtained by shuffling trials before Granger causality computation. Second, there had to be an asymmetry of directional coupling when subtracting Granger causality from A to B and B to A, i.e., their difference had to be significantly different from zero as assessed by two-tailed t-tests across electrode pairs (p < 0.05 FDR corrected). Third, when contrasting results from real (i.e., time-forward) and time-reversed data, the sign of the interaction had to flip. We used this conservative approach to preclude spurious interactions due to, e.g., volume conduction effects^25^. If all three criteria were fulfilled, a given functional connection between areas A and B was considered as valid in the sense that it statistically exhibited true interaction.

This assessment of Granger causality also lay the grounds for all further analyses: for window differences, we subtracted Granger causality of a given pair of temporal windows that were the same as those used for spiking information analysis (fixation, stimulus presentation, early and late delay periods), and for condition differences, we subtracted Granger causality from a given pair of conditions for each of those temporal windows. Window and condition differences were assessed for significance by two-tailed t-tests in addition to fulfilling the above criteria. As for neural information analyses, alpha levels for Granger causality statistics were set to 0.05 and p-values were FDR-corrected to account for multiple comparison.

All data analyses were performed in MATLAB using custom scripts and the FieldTrip^80^ toolbox.

## Supplemental Information

**Figure S1:**
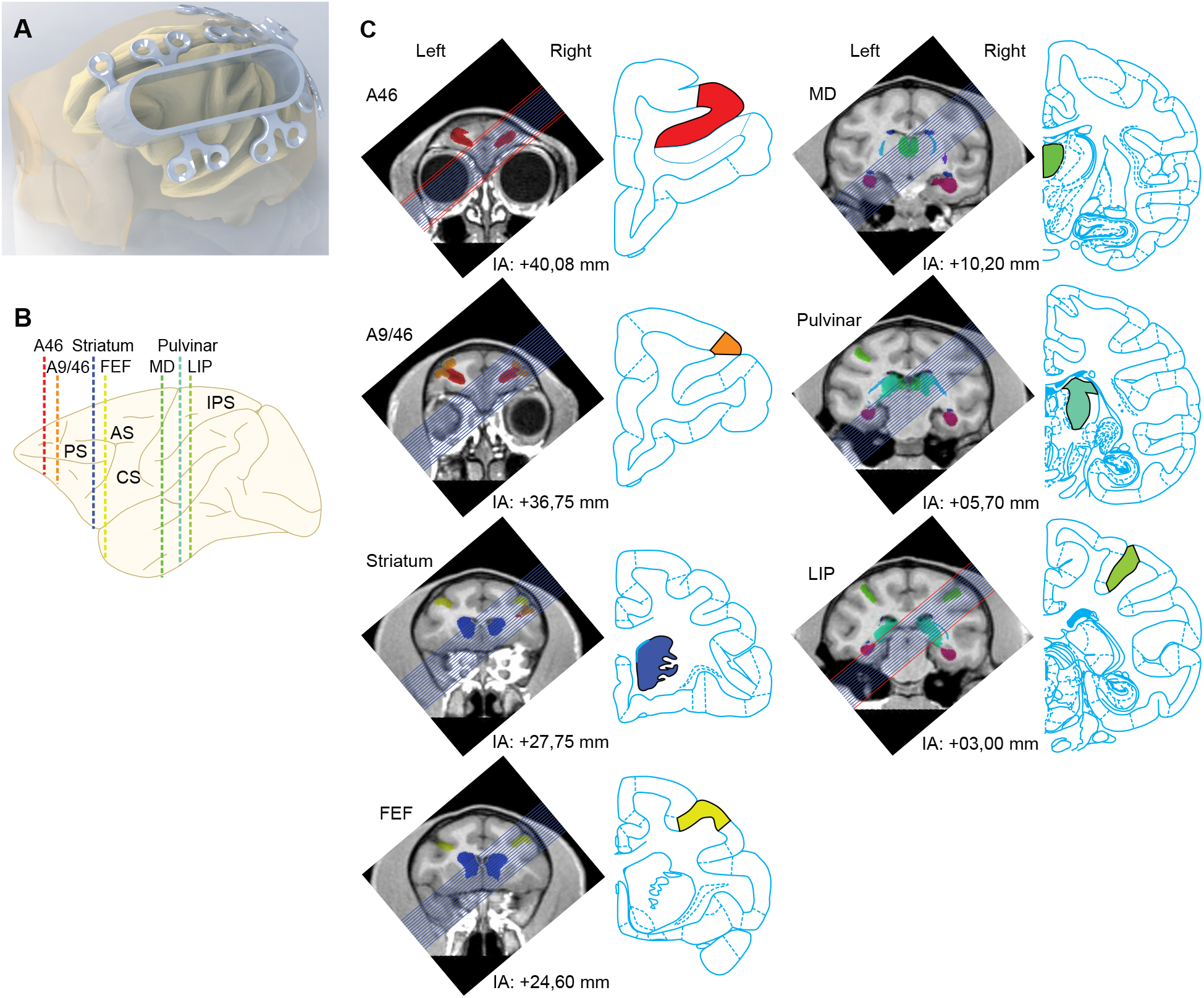
Recording approach. We used custom-designed titanium chambers spanning large parts of the anterior-posterior (AP) extent of one hemisphere, custom-designed microdrives and grids to record broadband neuronal activity from multiple cortical and subcortical structures simultaneously. (**A**) 3D-rendering of the titanium chamber model and of the headpost-base on the skull and brain of one of the monkeys. (**B**) Lateral view of a monkey brain from the standard monkey brain atlas of Paxinos and Watson illustrating the position of frontal sections used in (C) and corresponding most closely to the AP positions of the probes used in a representative recording session from one monkey. Colors correspond to those used for ROI illustrations in (C). (**C**) We mapped electrode trajectories (blue lines) onto the individual animal’s structural MRI including colored regions of interest (ROI). Frontal MRI sections from one monkey and most closely matching frontal sections from the standard monkey brain atlas of Paxinos and Watson. AP coordinates are given with reference to interaural (IA) zero. Colored ROIs indicate target areas. We rotated MRI images by 50 degrees to account for the chamber’s inclination and to ease comparison with the standard atlas sections. Red lines indicate trajectories blocked by the curved parts of the chamber.

